# The ground truth of the Data-Iceberg: Correct Meta-data

**DOI:** 10.1101/2021.12.17.473021

**Authors:** Aylin Caliskan, Seema Dangwal, Thomas Dandekar

## Abstract

Biological molecular data such as sequence information increase so rapidly that detailed metadata, describing the process and conditions of data collection as well as proper labelling and typing of the data become ever more important to avoid mistakes and erroneous labeling. Starting from a striking example of wrong labelling of patient data recently published in Nature, we advocate measures to improve software metadata and controls in a timely manner to not rapidly loose quality in the ever-growing data flood.

## Identical data with different labels

Recently, Han et al. (2021) reported “*Identification of SARS-CoV-2 inhibitors using lung and colonic organoids*”. Using alveolar type-II-like cells permissive to SARS-CoV-2 infection the authors performed a high-throughput screen of approved drugs to identify entry inhibitors of SARS-CoV-2 compared to uninfected controls, such as imatinib, mycophenolic acid and quinacrine dihydrochloride.

We intended to study whether or not the alveolar epithelial cells react to SARS-CoV-2, and therefore we compared these data to those in the article “*Imbalanced Host Response to SARS-CoV-2 Drives Development of COVID-19*” by Blanco-Melo et al. published in cell (2020). These authors revealed a unique and inappropriate inflammatory response, defined by low levels of type I and III interferons juxtaposed to elevated chemokines and high expression of IL-6 compared to controls. To our surprise, we learned that the Meta-data were entirely wrong annotated (see extended data):

We found irregularities in RNA-Seq data^1^ (Fig. 1, Han *et al*.): two of their human lung tissue samples (GSM4697983 and GSM4697984 from GSE155241) appear to be exactly the same as two other unrelated samples of human lung tissue that were generated during research for the *Cell* publication^4^.

**Fig 1.**
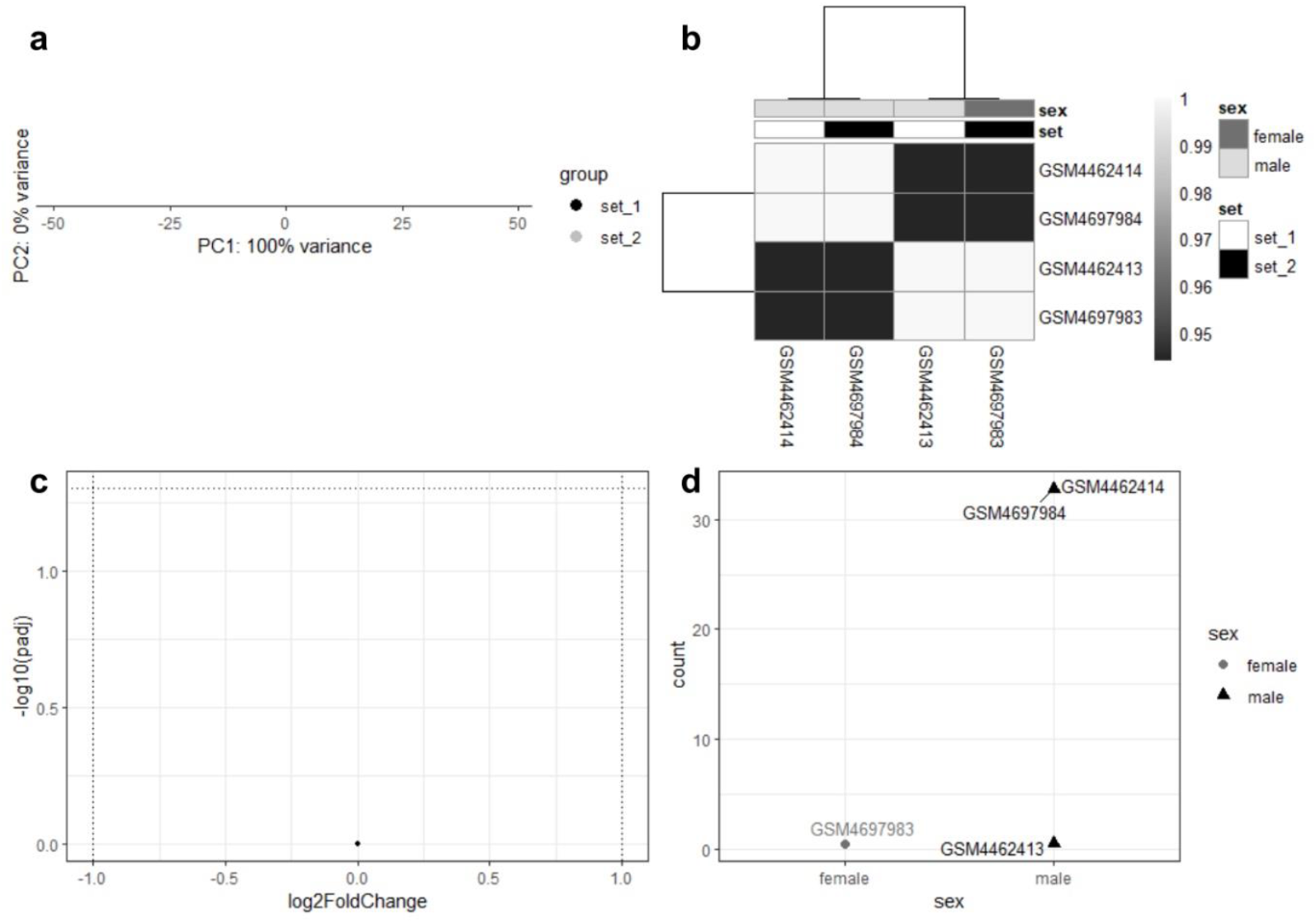
The figure shows the similarities between the samples using several standard analysis methods in R. **a**, Principal component analysis (PCA) comparing the two datasets. Set_1 contains the two healthy human lung tissue samples GSM4462413 and GSM4462414 from GSE147507 by Blanco-Melo et al.^2^. Set_2 contains two of the three healthy human lung tissue samples analysed by Han et al.^1^ (GSM4697983 and GSM4697984 from GSE155241). **b**, A heat map comparing the four samples with each other. The information bars at the top indicate the sex of the donor and to which dataset the samples belong. Both clustering and the colours in the heat map indicate the similarity of the samples. **c**, A volcano plot usually visualises differentially expressed genes (DEGs) in a shape that reminds of a volcano. Using the data of Set_1 compared to the data of Set_2 to generate a volcano plot results in a single dot, which indicates that there are no differentially expressed genes and that the expressed genes of both datasets and their expression are exactly the same. **d**, Gene count comparison using Xist. Xist is responsible for the dosage equivalence of X-linked genes in males and females and the inactivation of the second X-chromosome in females. As its expression is linked to the inactivated X-chromosome, it is expected to be typically expressed in females^3^. Therefore, the samples were grouped according to the recorded sex (metadata / personal communication). The count numbers of Xist expression of GSM4462413 and GSM4697983 as well as the Xist expression of GSM4697984 and GSM4462414 are exactly the same, which also indicates the similarity of the respective samples.

Sample GSM4462413, from a healthy 77-year-old-man in Blanco-Melo’s study^2^ (SRA metadata), appears to be the same as GSM4697983 (Han et al)^1^ labelled as healthy 56-year-old-woman. In addition, sample GSM4462414, from a healthy 72-year-old male in Blanco-Melo^2^ (SRA metadata), expresses gene sets and counts similar to the sample GSM4697984 from Han’s study^1^, generated from a healthy 66-year-old-man’s bio specimen (personal communication).

In other words, the results of both high impact factor studies are in doubt, as the control samples are clearly mixed-up and hence we do not really know how SARS-CoV2 infected cells behave compared to the control cells and who used the correct controls.

Is this a rare incident? No, mistakes with metadata happen very often, each time data are carelessly stored and meta-data (experimental conditions, samples, numbers, clinical features) either not entered or are mislabelled. However, we can pinpoint mistakes as the one above only rarely, as this requires close comparisons of the control data sets in unrelated publications, or, to be more general, basic and more refined quality checks on transcriptome data-sets and other omics data-sets across all public data-sets. We give in the following a brief overview on such errors to stress the necessity to do such checks.

## Metadata errors happen even more often

### Nucleotide Annotation errors

The notion of abounding mistakes in data and metadata is correct as you can already see with basic mistakes regarding nucleotide errors (Park et al., 2021):

Park et al. developed a semi-automated screening tool Seek & Blastn and, alarmingly, found 712 papers with wrongly identified sequences in five corpora (two journals and three targeted corpora)^4^. According to Google Scholar, these papers were cited 17,183 times in March 2021, including clinical trials^4^.

At the time of publication, up to 4% of the problematic papers in each corpus had already been cited at least once by clinical trials. However, Park and colleagues also analysed the papers further. They found a high probability of 15-35% for the problematic publications in each of the five corpora, respectively, to be cited in clinical research in future as they resemble papers that have been cited in clinical research^4^. Hence, there is serious concern that about a quarter of these publications are likely to impair clinical research by misinforming or distracting the development of potential cures.^4^

### Transcriptome meta-data errors

Mishandling of meta-data is always possible for transcriptome data sets, as well as for all other omics data-sets (proteome, phosphor-proteome, metabolome, genomics) and, the problem is, wrong metadata, wrong labels, wrong conditions, mislabelled controls are very difficult to spot and correct in retrospect. The control best is done using all info present at submission. The big wave of wrong annotation and big data errors is steadily rising:

The sheer amount of data and its rapid growth further complicate finding such errors. Thus, almost in 2013, the amount of publicly accessible gene-expression data sets was about to hit the one-million-milestone^6^. Using these data was starting to get recognised as a valid method of gathering research data, as 20% of the data sets that were deposited in 2005 had been cited by the year 2010^6^. Additionally, 17% of the data sets deposited in 2007 had been cited by the end of 2010^6^, underlining the increasing importance of the GEO database. By the end of 2020, GEO entries reached 4 Mio and only a year later, we are at 4,713,471 samples (November 2021). Data rise leads to error rise.

### Many annotation errors regarding sex

Toker et al.^5^ used a human transcriptomics studies to compare the sex of the subjects that was annotated in the metadata with what they termed the “gene-sex”, the sex of the subject determined by analysing the expression of the female-specific gene XIST and the male-specific genes KDM5D and RPS4Y1 ^5^. Their analysis revealed that 46% of the 70 datasets examined in the study were mislabelled^5^. Thus, they had a closer look at the respective original studies (n=29) and tried to find out whether the incorrect annotation was due to a miscommunication while uploading the data on the GEO database. Moreover, 12 of these 29 studies provided enough information in the publication to show, alarmingly, that the discrepant sex labels had already been present in the publication^5^.

Finally, Toker and colleagues compared four datasets that used samples from the same collection of subjects. Out of these four datasets, two contained mismatched samples^5^. Notably, the mismatched samples differed between the two datasets, and the respective samples were correctly annotated in the other two studies, indicating that the samples had been mislabelled instead of an error during the recording of the subject’s sex^5^.

## Discussion

The conclusions of both high-ranking papers in our start example are in doubt particular due to their unclear annotated controls. These data have to be rechecked carefully, but as a general rule, also the repositories should have automatic checking routines for such mistakes. This is easy to achieve: if every entry is checked for novelty of the raw data or even the partial overlap with the stored data, it is easy to identify this type of mistakes.

Based on the studies above, it is reasonable to assume that individual metadata errors are normal distributed and that for a low percentage of publications (at least 1-3%) a fatal binary error occurs such as wrong sex or mix-up of control and treatment while a substantial larger fraction (5x more?) has then minor quality issues.

Unfortunately, not all labelling errors can be found as easily as the wrong sex in the metadata. Most errors regarding the metadata are very difficult to spot in retrospect. Samples that erroneously got labelled as control samples are harder to identify and might cause even more damage to research, especially if there is only a limited number of samples with the particular condition available.

Big data are constantly increasing and this inevitably increases the work load on the people having to handle them. Unless there are automatic quality controls, cross-comparisons to validate metadata, checks and counter-checks that the entered information is correct we are to drown in errors as the number of personnel involved in databanks is certainly not increasing at the same pace as data generation. There is the well-known reproducibility crisis^7^ triggered by a bias in publishing positive results and not showing negative results or even omitting most of the confounding data. There are the correct reservations of statisticians regarding statistical biases, too small samples, extravagant claims and missing controls^8,9^. However, from the start of any scientific study, good data need good curation. If there is no exponential increase in automatic data and meta-data quality controls, we will experience a steady decline in data quality, inverse proportional to data growth.

## Supporting information

we learned that the Meta-data were entirely wrong annotated (see extended data …

## References

1. Han, Y. et al. Identification of SARS-CoV-2 inhibitors using lung and colonic organoids. Nature 589, 270–275 (2021). doi: https://doi.org/10.1038/s41586-020-2901-9

2. Blanco-Melo, D. et al., Imbalanced Host Response to SARS-CoV-2 Drives Development of COVID-19, Cell 181, 1036–1045.e9 (2020). doi: https://doi.org/10.1016/j.cell.2020.04.026

3. Brockdorff, N. et al. Conservation of position and exclusive expression of mouse Xist from the inactive X chromosome. Nature, 351(6324):329–31 (1991). doi: 10.1038/351329a0. PMID: 2034279.

4. Park, Y., et al. (2021) Human gene function publications that describe wrongly identified nucleotide sequence reagents are unacceptably frequent within the genetics literature, bioRxiv 2021.07.29.453321; doi: https://doi.org/10.1101/2021.07.29.453321

5. Toker L, Feng M and Pavlidis P. Whose sample is it anywayã Widespread misannotation of samples in transcriptomics studies [version 2; peer review: 2 approved, 1 approved with reservations]. F1000Research 5:2103 (2016) (https://doi.org/10.12688/f1000research.9471.2)

6. Baker, M., Gene data to hit milestone [published correction appears in Nature.2012 Aug 2;488(7409):19]. Nature 487(7407):282–283 (2012). doi:10.1038/487282a

7. Baker, M. 1,500 scientists lift the lid on reproducibility. Nature 533, 452–454 (2016). https://doi.org/10.1038/533452a

8. Ioannidis, John P A. Why most published research findings are false. PLoS medicine vol. 2, 8, e124 (2005). doi:10.1371/journal.pmed.0020124

9. Ioannidis JP, Munafò MR, Fusar-Poli P, Nosek BA, David SP. Publication and other reporting biases in cognitive sciences: detection, prevalence, and prevention. Trends Cogn Sci. 18(5):235–241 (2014). doi:10.1016/j.tics.2014.02.010

