## Supplementary material for "The ground truth of the Data-Iceberg: Correct Meta-data": we learned that the Meta-data were entirely wrong annotated (see extended data …

### Additional Information on the Respective Datasets

The RNA-Seq data we intended to use in our study was derived from publicly available RNA-Seq data in the GEO database[10]

One GEO dataset was produced as part of the *Nature* publication "Identification of SARS-CoV-2 inhibitors using lung and colonic organoids" by Han *et al.* (GSE155241)[1]. The article was first published in October 2020 and has been accessed at least 45,000 times and cited 86 times in Web of Science and 116 times in CrossRef, according to *Nature*'s article metrics (as of the end of November 2021)[11].

The other GEO dataset was generated during research for the paper "Imbalanced Host Response to SARS-CoV-2 Drives Development of COVID-19", written by Blanco-Melo *et al.* (GSE147507)[2]. The authors published their research in May 2020 in *Cell* and have been cited over a thousand times (according to PlumX metrics 1,023 times in CrossRef, 1,364 times in Scopus, and 1,348 times in PubMed Central, accessed at the end of November 2021)[12].

Two samples of the GEO dataset by Han *et al.* (GSM4697983 and GSM4697984 from GSE155241)[1] appear to be duplicates or doppelgangers of two other, unrelated samples of the GEO dataset by Blanco-Melo *et al.* (GSM4462413 and GSM4462414 from GSE147507)[2].

All of these samples are said to be samples of healthy lung tissue that were used as controls during COVID research[1,2].
According to the respective publications, the tissue samples of human lungs that were not infected with SARS-CoV-2 were provided by different institutions and belonged to different donors[1,2].

### Information on the Respective Samples

Blanco-Melo *et al.* received their uninfected human lung samples via the Mount Sinai Institutional Biorepository and Molecular Pathology Shared Resource Facility (SRF) in the Department of Pathology[2]. According to their publication, all lung tissue samples were derived from over 60-year-old males[2]. The metadata, which is available via the SRA Run Selector, gives even more detail about the origin of the samples.

1. "Series15_HealthyLungBiopsy_1" (GSM4462413) was used as a healthy negative control and derived from a 77-year-old male, who received no treatment.
2. The donor of the second control sample, "Series15_HealthyLungBiopsy_2" (GSM4462414), was a healthy 72-year-old male.

Both samples were released on 2020-04-10 and sequencing data obtained via Illumina NextSeq 500 sequencing.

The Han *et al.* publication compared SARS-CoV-2 infected lung tissue and uninfected lung tissue[1]. Their human tissue samples were provided by the Weill Cornell Medicine Department of Pathology[1].

According to their supplementary information, they received three samples of uninfected lung tissue[1]:

1. “Healthy_1” (56, female),
2. “Healthy_2” (66, male) and
3. “Healthy_3” (38, female).

The release date of the samples is 2020-08-12, and the sequencing data were obtained using Illumina NovaSeq 6000.

The metadata of the files in the SRA run selector did not contain further information about the patients’ age and sex, and the information at GEO was ambiguous with:

1. GSM4697982 Healthy Lung autopsy [Healthy-lung]
2. GSM4697983 Healthy Lung autopsy_1 [HealthyLungBiopsy_1]
3. GSM4697984 Healthy Lung autopsy_2 [HealthyLungBiopsy_2]

Thus, we contacted the corresponding author to confirm whether the annotation we assumed was correct and received the following patient information (personal correspondence, July 2021):

1. GSM4697982 Healthy Lung autopsy [Healthy-lung] 38 Female
2. GSM4697983 Healthy Lung autopsy_1 [HealthyLungBiopsy_1] 56 Female
3. GSM4697984 Healthy Lung autopsy_2 [HealthyLungBiopsy_2] 66 Male

**Analyzing the Data**

Analyzing the data of both studies in R (version 4.1.0 (2021-05-18))[13], using R studio (version 1.4.1717, “Juliet Rose” (df86b69e, 2021-05-24) for Windows) and several packages (see the Materials and Methods for further information) revealed that GSM4462413 and GSM4697983 as well as GSM4462414 and GSM4697984 show an identical count and sequencing read distribution (see Figure 1 in the main text).

**Materials and Methods**

Data

Control samples of research regarding SARS-CoV-2 infection:

- Regarding the transcriptional response to SARS-CoV-2 infection, for the Blanco-Melo et al. publication “Imbalanced Host Response to SARS-CoV-2 Drives Development of COVID-19” (doi: https://doi.org/10.1016/j.cell.2020.04.026, GSE147507 (GSM4462413, GSM4462414)[2]
- Regarding Identification of Drugs Blocking SARS-CoV-2 Infection using hPSC-derived Lung and Colonic Organoids, for the Han at al. Publication “Identification of SARS-CoV-2 inhibitors using lung and colonic organoids” doi: https://doi.org/10.1038/s41586-020-2901-9, GSE155241 (GSM4697983, GSM4697984)[1]).

Hardware

The downloads and the subsequent preparation steps were performed on a PC with AMD Ryzen 9 3900X, 12-Core Processor, 64.0 GB RAM, 64-Bit-Operating System, and an x64-based processor.

RNA-Seq Data Preparation

While data download and preparation were done in a virtual Ubuntu environment (Ubuntu 20.04.2 LTS (OS-Type: 64-bit) on a virtual machine (Virtual Box 6.1)), data analysis was performed using RStudio (version 1.4.1717, "Juliet Rose" (df86b69e, 2021-05-24) for Windows) on Windows 10 with R version 4.1.0 (2021-05-18)[13].

The RNA-Seq data were obtained by selecting the respective entries in the NCBI SRA Run Selector, downloading the SraRunTable (see Data S1) containing the metadata, and downloading the SRA files via prefetch (version 2.11.0) and fastq-dump (version 2.11.0). All files were converted into fastq format using the NCBI SRA Toolkit[14]. FastQC (version 0.11.9)[15] and multiqc (version 1.11)[16] were used for quality control (in several conda environments (conda version 4.10.3).

After STAR alignment (version 2.7.9a)[17], using the comprehensive gene annotation PRI and the genome sequence, primary assembly (GRCh38) PRI of Gencode version 37 [18] and transcript quantification with RSEM (version 1.3.3)[19], the resulting files were analyzed in RStudio.

Data Analysis in RStudio

We compared the four samples in RStudio, using the following packages: ggplotify (version 0.0.7)[20], ggrepel (version 0.9.1)[21], ggplot2 (version 3.3.4)[22], cowplot (version 1.1.1)[23], RColorBrewer (version 1.1-2)[24], pheatmap (version 1.0.12)[25], tidyverse (version 1.3.1)[26], DESeq2 (version 1.32.0)[27] with apeglm[28], tximport (version 1.20.0)[29], BiocManager (version 1.30.16)[30].

**References**

10. R. Edgar, M. Domrachev, A. E. Lash, Gene Expression Omnibus: NCBI gene expression and hybridization array data repository, *Nucleic Acids Research* **30**, 207–210 (2002). doi: https://doi.org/10.1093/nar/30.1.207

11. Nature article metrics (2021), (available at: https://www.nature.com/articles/s41586-020-2901-9/metrics)

12. PlumX Metrics (2021), (available at: https://plu.mx/plum/a/?doi=10.1016/j.cell.2020.04.026&theme=plum-jbs-theme&hideUsage=true )

13. R Core Team, R: A language and environment for statistical computing. R Foundation for Statistical Computing, Vienna, Austria (2021). URL https://www.R-project.org/

14. D.L. Wheeler, T. Barrett, D.A. Benson, *et al.* Database resources of the National Center for Biotechnology Information. *Nucleic Acids Res*. **34** (Database issue):D173-D180 (2006). doi:10.1093/nar/gkj158, available at: http://ncbi.github.io/sra-tools/

15. S. Andrews, FastQC: a quality control tool for high throughput sequence data. (2010). Available online at: http://www.bioinformatics.babraham.ac.uk/projects/fastqc

16. P. Ewels, M. Magnusson, S. Lundin, M. Käller, MultiQC: Summarize analysis results for multiple tools and samples in a single report. *Bioinformatics* (2016). doi: 10.1093/bioinformatics/btw354 PMID: 27312411
17. A. Dobin, C.A. Davis, F. Schlesinger, *et al.* STAR: ultrafast universal RNA-seq aligner. *Bioinformatics* **29**(1):15-21 (2013). doi:10.1093/bioinformatics/bts635
18. A. Frankish, M. Diekhans, A.M. Ferreira, *et al.* GENCODE reference annotation for the human and mouse genomes. *Nucleic Acids Res*. **47**(D1):D766-D773 (2019).
19. B. Li, C.N. Dewey, RSEM: accurate transcript quantification from RNA-Seq data with or without a reference genome. *BMC Bioinformatics* **12,** 323 (2011).
20. Guangchuang Yu (2021). ggplotify: Convert Plot to 'grob' or 'ggplot' Object. R package version 0.0.7. https://CRAN.R-project.org/package=ggplotify
21. K. Slowikowski (2021). ggrepel: Automatically Position Non-Overlapping Text Labels with 'ggplot2'. R package version 0.9.1. https://CRAN.R-project.org/package=ggrepel
22. H. Wickham. ggplot2: Elegant Graphics for Data Analysis. *Springer-Verlag New York*, (2016).
23. C.O. Wilke (2020). cowplot: Streamlined Plot Theme and Plot Annotations for 'ggplot2'. R package version 1.1.1. https://CRAN.R-project.org/package=cowplot
24. E. Neuwirth (2014). RColorBrewer: ColorBrewer Palettes. R package version 1.1-2. https://CRAN.R-project.org/package=RColorBrewer
25. Raivo Kolde (2019). pheatmap: Pretty Heatmaps. R package version 1.0.12. https://CRAN.R-project.org/package=pheatmap
26. Wickham et al., (2019). Welcome to the tidyverse. Journal of Open Source Software, 4(43), 1686, https://doi.org/10.21105/joss.01686
27. M.I. Love, W. Huber, S. Anders, Moderated estimation of fold change and dispersion for RNA-seq data with DESeq2 *Genome Biology* **15**(12):550 (2014)
28. A. Zhu, J.G Ibrahim, M.I. Love, Heavy-tailed prior distributions for sequence count data: removing the noise and preserving large differences. *Bioinformatics* (2018). https://doi.org/10.1093/bioinformatics/bty895
29. C. Soneson, M.I. Love, M.D. Robinson: Differential analyses for RNA-seq: transcript-level estimates improve gene-level inferences. F1000Research (2015)
30. M. Morgan (2021). BiocManager: Access the Bioconductor Project Package Repository. R package version 1.30.16. https://CRAN.R-project.org/package=BiocManager
